# Identification of annotation artifacts concerning the *CHALCONE SYNTHASE* (CHS)

**DOI:** 10.1101/2023.03.18.533251

**Authors:** Martin Bartas, Adriana Volna, Jiri Cerven, Boas Pucker

## Abstract

**Objective:** Chalcone synthase (CHS) catalyzes the initial step of the flavonoid biosynthesis. The CHS encoding gene is well studied in numerous plant species. Rapidly growing sequence databases contain hundreds of CHS entries that are the result of automatic annotation. In this study, we evaluated apparent multiplication of CHS domains in *CHS* gene models of four plant species.

**Main findings:** *CHS* genes with an apparent triplication of the CHS domain encoding part were discovered through database searches. Such genes were found in *Macadamia integrifolia, Musa balbisiana, Musa troglodytarum*, and *Nymphaea colorata*. A manual inspection of the *CHS* gene models in these four species with massive RNA-seq data suggests that these gene models are the result of artificial fusions in the annotation process. While there are hundreds of apparently correct CHS records in the databases, it is not clear why these annotation artifacts appeared.

## Introduction

Flavonoids are one of the most important groups of specialized plant metabolites. Their enormous chemical diversity results in a plethora of biological functions [1]. Most noticeable are the anthocyanins which can provide blue to red coloration to flowers. Flavonoids are distributed across the whole plant kingdom. Given the visual phenotype of several flavonoids, this pathway was established as a model system for plant metabolism and transcriptional regulation [2, 3]. Today, the flavonoid biosynthesis and its regulation are among the best studied processes in plants. Numerous studies are published every year which investigate the flavonoid biosynthesis and the corresponding regulators in different plant species.

One of the best studied enzymes in the flavonoid biosynthesis pathway is the chalcone synthase (CHS). This enzyme catalyzes the first committed step of the flavonoid biosynthesis, particularly the reaction leading to the formation of the naringenin chalcone from one oumaroyl-CoA and three malonyl-CoA [4]. CHS belongs to the type III polyketide synthases, a larger protein family that also harbors several closely related enzymes like the stilbene synthase (STS) [5]. CHS and STS differ by only two functionally important residues which are Q166 and Q167 in the *Arabidopsis thaliana* CHS sequence [6]. The gene structure of *CHS* comprises usually two coding exons [7].

Numerous plant genomes are sequenced every year [8] and the rapid development of long read sequencing technologies enables the generation of high quality genome sequences [9]. Since an automatic annotation of all genes in the flavonoid biosynthesis and many regulators is possible [10, 11], it is feasible to perform large scale analyses of gene families acting in this pathway. Analyses at this scale are inherently prone to errors concerning individual species and sequences that require human intervention at the final interpretation step. A systematic analysis of domains in the chalcone synthase revealed an apparent triplication in several species.

Here, we describe an investigation of several *CHS* gene models that appeared to encode a protein with a triplicated CHS domain. A manual inspection of multiple cases based on transcriptomic data suggests mis-annotation leading to artificially fused gene models.

## Methods

### Identification of sequences with potential CHS domain duplications

A BLASTp analysis with the *Nymphae colorata* CHS (XP_049936683.1) as query and nr (containing 545546009 sequences) as subject was performed through the NCBI webportal (https://blast.ncbi.nlm.nih.gov/Blast.cgi?PAGE=Proteins) was performed on the 2nd of June 2023. The word size was set to 5 and an e-value cutoff of 10^−30^ was applied. Matched sequences longer than 500 amino acids were manually screened for CHS domain duplications or triplications. Redundant sequences were removed from the list.

### Evaluation of gene models via RNA-seq read mapping

The genome sequences and corresponding annotations of four species with apparent fusion of multiple CHS gene copies were selected for manual inspection: *Musa balbisiana* [12], *Musa troglodytarum* [13], *Macadamia integrifolia* [14], and *Nymphaea colorata* [15]. All available paired-end RNA-seq data sets of these species were retrieved from the Sequence Read Archive via fastq-dump [16] (Additional File 1 in [17]). Read pairs were aligned to the genome sequence with STAR v2.5.1b [18, 19] in 2-pass-mode using a minimal similarity of 95% and a minimal alignment length of 90% as previously described [20]. The resulting BAM files were processed with customized Python scripts [17]. All reads mapped to the locus of interest were extracted from each individual BAM file with samtools v1.15.1 [21]. These subset BAM files were merged to produce one BAM file per species that contains all reads belonging to the locus of interest (Additional File 2, Additional File 3, Additional File 4, Additional File 5 in [17]). The final BAM file was visualized with Integrative Genomics Viewer (IGV) [22].

### Analysis and visualization of RNA-seq read mapping coverage

A previously developed [23] Python script was applied to convert BAM files into coverage files that list the number of aligned reads per genomic position. These coverage files served as the basis for the generation of gene model focussed coverage plots [17]. These plots visualize the number of aligned reads per position around a given locus of interest. High coverage with RNA-seq reads indicates exon positions while introns are characterized by very low or no RNA-seq read coverage at all. Introns are not included in the final mRNAs and usually only mature mRNAs are extracted with standard RNA extraction protocols used for RNA-seq experiments.

### Comparison of sequence similarity

Sequences around the CHS loci of interest were retrieved from the respective genome sequence using the Python script seqex3.py v0.5 [24]. Sequence similarity between the different domains predicted in the inspected sequences was analyzed by constructing dot plots with dotplotter v0.1 [25] using a k-mer size of 31. Dotplotter compares two sequences by generating k-mers from one given sequence and identifying matches in the other sequence. The corresponding k-mer positions in both sequences are visualized in a dotplot. This allows an intuitive visualization of repeats.

## Results

An analysis of the CHS sequences retrieved from the NCBI revealed apparent fusion proteins harboring multiple (two or three) CHS domains in 28 protein sequences across different plant species (Additional File 6 in [17]). More specifically, 8 sequences showed a triplication of the CHS domain, while 20 sequences showed a duplication. Representative examples of predicted proteins with multiple CHS domains were identified in *Macadamia integrifolia, Musa balbisiana, Musa troglodytarum*, and *Nymphae colorata* (**Fig. 1**). A comparison of each *CHS* locus in the four species against itself revealed the expected high similarity (Additional File 7 in [17]).

**Fig. 1:**
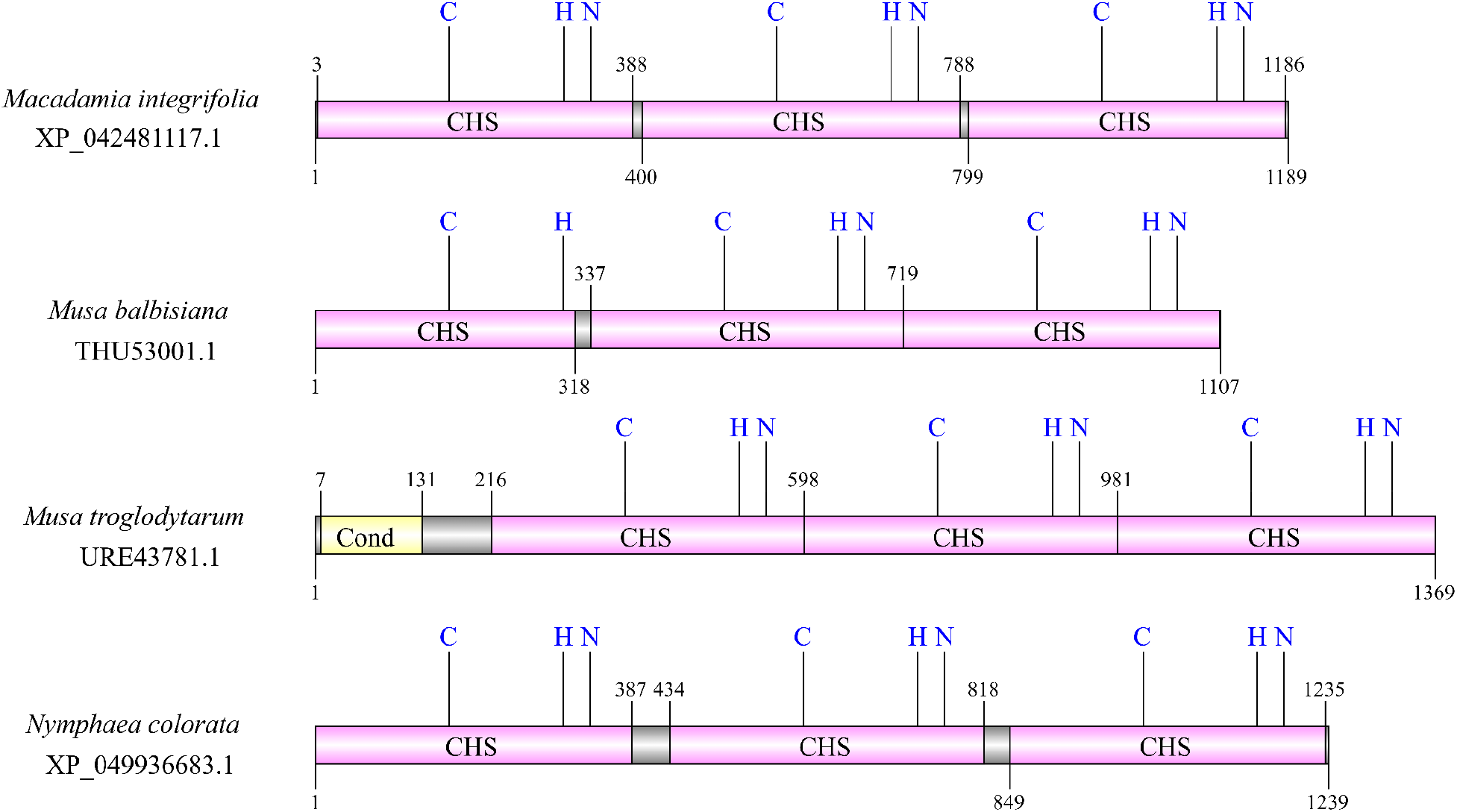
Domain composition of protein sequences with apparent “CHS triplication”. CHS abbreviation in figure stands for *Chalcone synthase domain*, and Cond abbreviation stands for *Chalcone and stilbene synthases, N-terminal domain*. So-called catalytic triad consisting of conserved Cysteine (C), Histidine (H), and Asparagine (N) amino acid residues is always depicted (if preserved).

Loci with an unexpected *CHS* annotation were inspected in an RNA-seq read mapping (**Fig. 2**). The annotation indicates a single *CHS* gene encoding three CHS domains, but the results of our RNA-seq read mapping suggest that there are multiple individual *CHS* genes. The coverage continuously drops towards the ends of several exons. That is an indication of the end of a gene. In contrast, an abrupt drop in coverage to almost zero would indicate a splice site. There is almost no coverage between some of the annotated exons and individual exons show very different RNA-seq coverage. There are also substantial fractions of annotated exons without RNA-seq support. It is not expected that exons at the 5’- and 3’-end of a gene have high coverage, while the enclosed exons have low coverage. The underlying RNA-seq read mapping files of *Macadamia integrifolia* (Additional File 2 in [17]), *Musa balbisiana* (Additional File 3 in [17]), *Musa troglodytarum* (Additional File 4 in [17]), and *Nymphae colorata* (Additional File 5 in [17]) are available for in depth inspection. Detailed instructions are provided to enable researchers to perform a similar investigation of other cases of potential annotation artifacts [17].

**Fig. 2:**
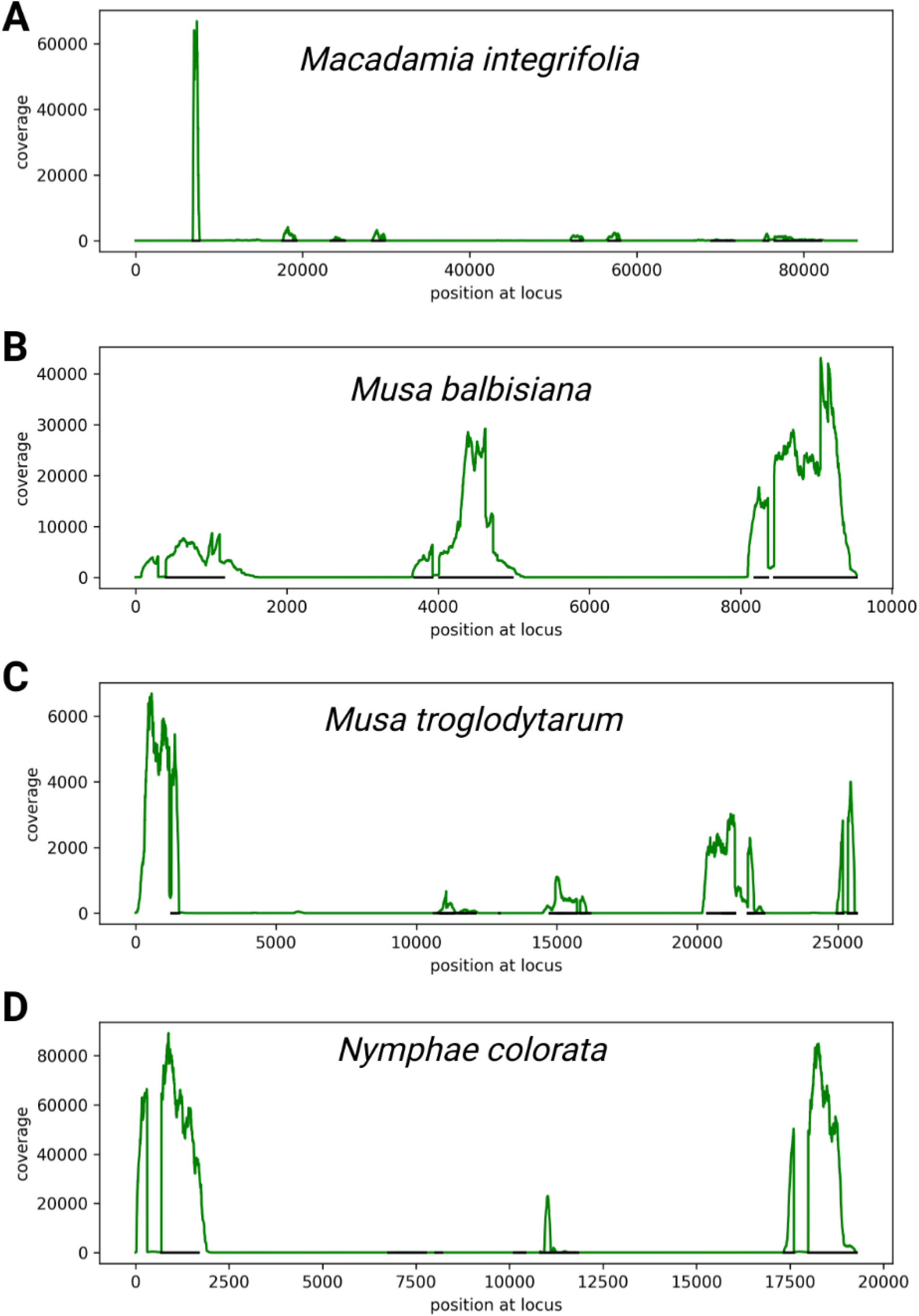
RNA-seq read coverage of *CHS* loci in *Macadamia integrifolia* (A), *Musa balbisiana* (B), *Musa troglodytarum* (C), and *Nymphae colorata* (D). The annotation suggests that a single gene is spanning the entire displayed locus, but the continuously dropping coverage towards the end of exons (black lines) in our RNA-seq read mapping suggests that there are multiple individual genes.

## Discussion

We outlined how surprising results of a quick database search and in-depth inspection of these entries resulted in the identification of most likely annotation artifacts. We investigated the assembly and annotation process underlying the annotations of the four analyzed species. The *Macadamia integrifolia* assembly was produced with MaSuRCA v3.2.6 and ALLMAPS Jul-2019 [14, 26, 27]. The *Musa balbisiana* and *M. troglodytarum* genome sequences were assembled with wtdbg v1.2.8 and NextDenovo v2.4.0, respectively [12, 13, 28, 29]. The *Nymphae colorata* assembly was produced by Canu v1.3 [15, 30]. All these assemblers are well established tools that have been deployed in numerous plant genome sequencing projects. The annotation was produced by Gnomon [31] which is the default annotation pipeline operated at the NCBI. There is no indication how any of these steps could have caused the mis-annotation. Gnomon was also applied on many other plant genome sequences and usually generated *CHS* annotations comprising only a single CHS domain [32–34]. The repetitive regions with multiple *CHS* copies in an array might contribute to the mis-annotation as suggested for repeats before [35].

To the best of our knowledge, there are no reports that validated a CHS protein with multiple repeated domains. Despite all efforts, it might not be possible to automatically maintain a perfect database free of any mis-annotations. Despite technological advances, manual inspection and correction might be required in some cases [36]. SWISS-PROT is an initiative to establish a collection of sequences that were curated by experts [37, 38]. However, such a manual inspection is not a scalable approach given the rapid growth of sequence collections due to improved sequencing technologies [8, 9]. As most records in the database are likely correct, users should carefully inspect any substantially deviating records. Especially sequences, which appear as exciting discoveries, should be considered as suspicious and thorough inspection is required.

Unexpected sequences should be carefully checked, because annotation artifacts are more likely than striking differences between closely related species. We describe our strategy for the gene model investigation in detail to enable repetition by others [17].

1. A comparison with orthologs in many other species is often a way to identify unexpected sequence properties. Sequences should only differ systematically if there are particularly striking events during evolution. If such events are not known, any major sequence differences could point to artifacts and should be considered as such.
2. Generally, the alignment of RNA-seq reads or other types of transcriptomic evidence should be used to gain insights into the structure of genes. It is important to use a proper split read aligner like STAR [18, 19] or HISAT2 [39] for this step. The coverage should be consistent or should drop continuously towards the ends of a gene. Splice sites within a gene are characterized by abrupt changes of the coverage to almost zero. In addition, introns should be spanned by a number of reads that is almost equivalent to the coverage at the border of the flanking exons.
3. If no suitable datasets are available, it is also possible to validate the gene structure and sequence through amplification via RT-PCR followed by Sanger sequencing [40]. As this approach is more time consuming and requires more financial resources, the aforementioned data-reuse approaches should be applied first.

## Limitations

Our investigation was restricted to four examples of apparent CHS domain triplication events within a single *CHS* gene in four different plant species. The gene models in question were contradicted by the analyzed RNA-seq datasets. However, we cannot rule out the possibility that such modified CHS versions exist elsewhere.

## Abbreviations

CHS: Chalcone synthase
STS: Stilbene synthase
ER: Endoplasmic reticulum
IGV: Integrative Genomics Viewer

## Declarations

### Ethics approval and consent to participate

Not applicable

### Consent for publication

Not applicable

### Availability of data and materials

The developed scripts, a detailed description for the inspection of suspicious gene models, and additional sequence collections are available via GitHub: https://github.com/bpucker/CHS. All analyzed data sets are publicly available (Additional File 1 in [17]). Genome sequences and corresponding annotations were retrieved from the BananaGenomeHub (https://banana-genome-hub.southgreen.fr/) and the NCBI (https://www.ncbi.nlm.nih.gov/), respectively.

### Authors’ contributions

MB and BP planned the study. BP wrote the software and performed the bioinformatic analysis. MB, AV, JC and BP interpreted the results, and wrote the manuscript. All authors have read the final version of the manuscript and approved its submission.

## Acknowledgements

This work was supported by the BMBF-funded de.NBI Cloud within the German Network for Bioinformatics Infrastructure (de.NBI) (031A532B, 031A533A, 031A533B, 031A534A, 031A535A, 031A537A, 031A537B, 031A537C, 031A537D, 031A538A). Many thanks to the Bioinformatics Resource Facility (BRF) at the Center for Biotechnology (CeBiTec) at Bielefeld University for providing an environment to perform the computational analyses.

