## Supplementary figures and images for "Identification of annotation artifacts concerning the *CHALCONE SYNTHASE* (CHS)"

### Additional File 7

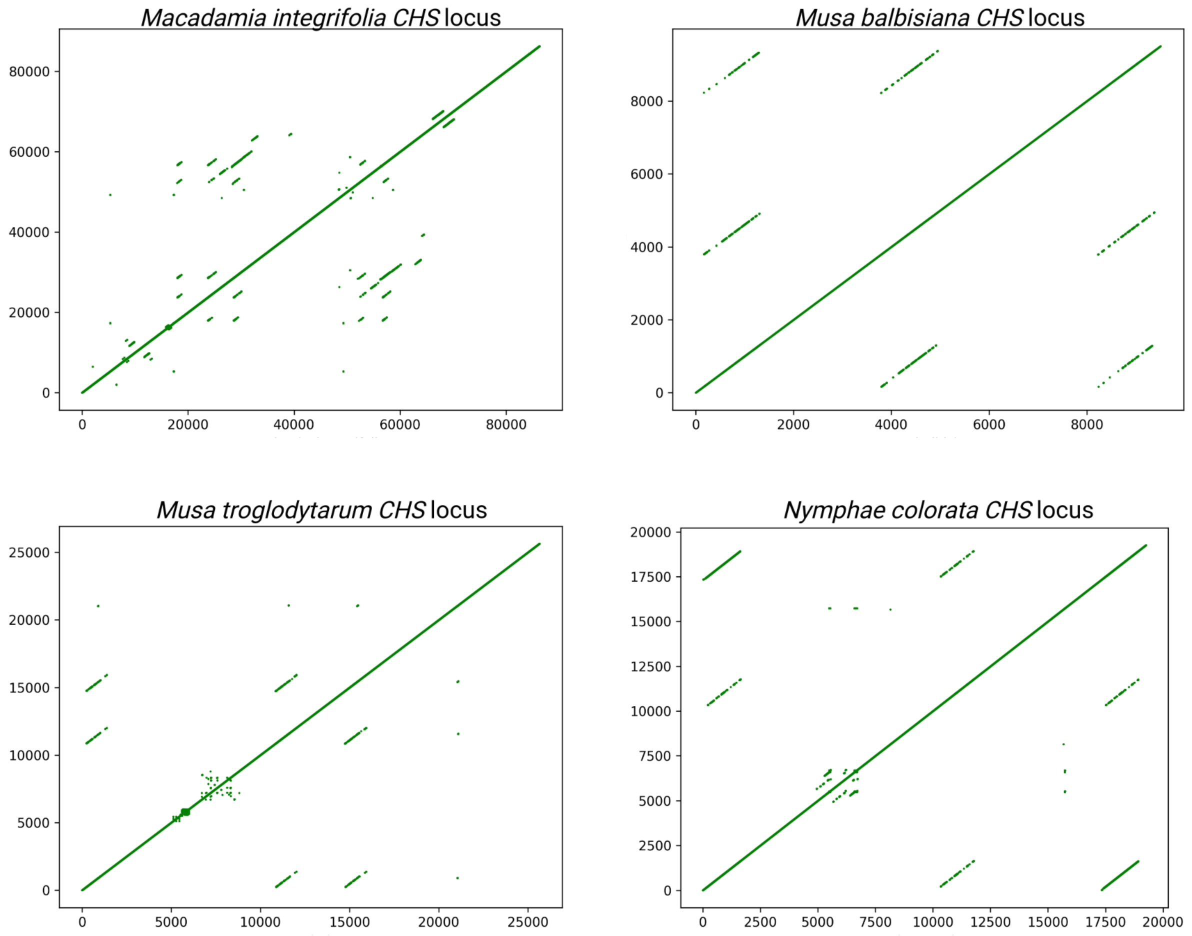
